# General Capillary Endothelial Cells Undergo Reprogramming into Arterial Endothelial Cells in Pulmonary Hypertension through HIF-2α/Notch4 Pathway

**DOI:** 10.1101/2024.02.13.580227

**Authors:** Bin Liu, Dan Yi, Xiaomei Xia, Karina Ramirez, Hanqiu Zhao, Yanhong Cao, Ankit Tripathi, Ryan Dong, Anton Gao, Hongxu Ding, Shenfeng Qiu, Vladimir V. Kalinichenko, Michael B. Fallon, Zhiyu Dai

## Abstract

Pulmonary arterial hypertension (PAH) is characterized by a progressive increase of pulmonary vascular resistance and obliterative pulmonary vascular remodeling that result in right heart hypertrophy, failure, and premature death. The underlying mechanisms of loss of distal capillary endothelial cells (ECs) and obliterative vascular lesion formation remain unclear. Our recent single-cell RNA sequencing, spatial transcriptomics analysis, RNASCOPE, and immunostaining analysis showed that arterial ECs accumulation and loss of capillary ECs were evident in human PAH patients and pulmonary hypertension (PH) rodents. Pseudotime trajectory analysis of the single-cell RNA sequencing data suggest that lung capillary ECs transit to arterial ECs during the development of PH. Our study also identified CXCL12 as the marker for arterial ECs in PH. Capillary EC lineage tracing approach using capillary specific-Dre;Tdtomato reporter mice demonstrated that capillary ECs gave rise to arterial ECs during PH development. Genetic deletion of HIF-2a or pharmacological inhibition of Notch4 normalized the arterial programming in PH. In conclusion, our study demonstrates that capillary endothelium transits to arterial endothelium through the HIF-2a-Notch4 pathway during the development of PAH. Thus, targeting arterial EC transition might be a novel approach for treating PAH patients.

---

Pulmonary arterial hypertension (PAH) is a devastating vascular disease with loss of distal capillaries and accumulation of PA endothelial cells (PAECs). There is a paradoxical observation that both loss of capillaries and accumulation of arterial ECs are evident in the lungs of PAH patients. Recent studies employing single-cell RNA sequencing (scRNA-seq) analysis demonstrated that lung ECs contain arterial, venous, general capillary (gCap) ECs (*Gpihbp1, Plvap*), aerocytes or aCap (*Car4, Ednrb*)^1^. gCap ECs function as stem/progenitor cells in capillary homeostasis and repair^1^. However, signaling pathways critical for PAEC specification during PH development remain elusive.

We previously generated *Egln1*^*Tie2Cre*^ mice (CKO), a novel pulmonary hypertension (PH) mouse model with progressive obliterative vascular remodeling and right heart failure^2^. Recently, we performed scRNA-seq analysis on lungs of *Egln1*^*f/f*^ (WT) mice and CKO mice. Our scRNA-seq data showed a decrease in both gCap and aCap EC proportions, but an increase in arterial EC proportion in CKO mice, suggesting an enrichment of arterial ECs in PH (**Fig. A, B**). The arterial ECs enriched in CKO mice express high levels of *Cxcl12, Sparcl1, Hspg2* **(Fig. C)**. To determine the localization of the EC subpopulation alteration, we performed spatial transcriptomics using the Visium Spatial Gene Expression platform. We then integrated the scRNA-seq and Visium datasets using Seurat and found that in the WT lung, arterial ECs are primarily found in proximal regions and gCap ECs are detected in distal microvascular regions. In contrast, CKO lungs showed an increase in arterial ECs in the distal microvascular bed and a decrease in gCap EC and aCap EC signatures (**Fig. D**). We then observed that CKO mice exhibited increased numbers of *Cxcl12*^+^/CD31^+^ in the distal microvasculature compared to the WT using immunostaining and RNASCOPE analysis (**Fig. E**). Via leveraging the publicly available scRNA-seq dataset^3^, we found that idiopathic PAH (IPAH) patients ECs also exhibited ∼3 fold higher of arterial ECs (40.3%) compared to donor lungs (10.7%) (**Fig. F and G**). Further comparison of human and mouse arterial EC markers showed that there were 61 common genes, including *CXCL12, SOX17, DEPP1, EDN1, HEY1* etc.

**Figure.**
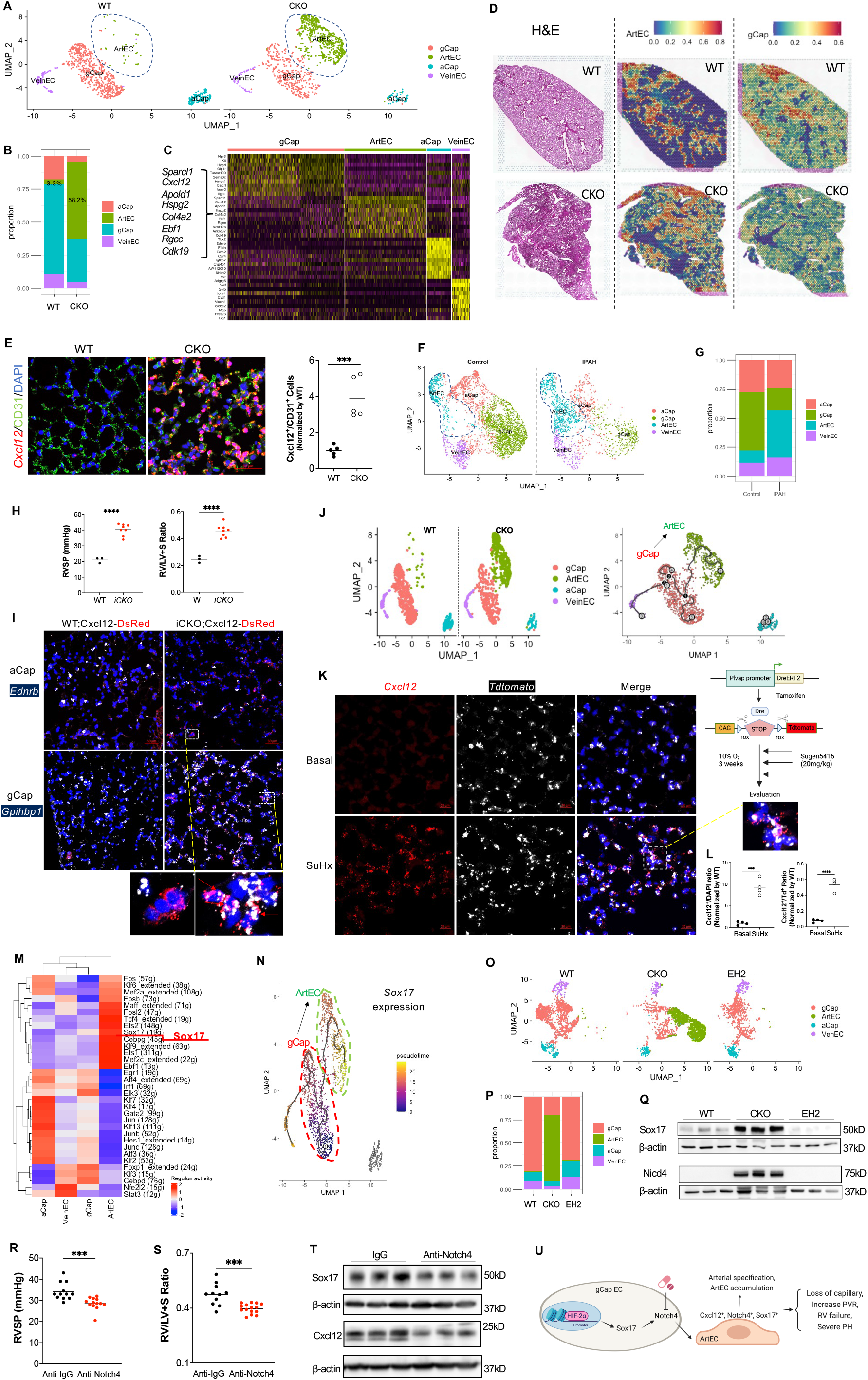
General Capillary Endothelial Cells Undergo Reprogramming into Arterial Endothelial Cells in Pulmonary Hypertension. **A and B**, A representative UMAP plot showing the abnormal EC subpopulations change with accumulation of arterial ECs (green) in both male and female CKO mice at the age of 3 months. **C**, A representative heatmap showing the top markers selectively enriched for each EC cluster, the data was generated by integrated data from WT and CKO. **D**, Integration of scRNA-seq and Visium data reveals increased arterial ECs and decreased gCap ECs in the distal capillary bed of CKO mice. The visualization shows the predicted spatial distribution of arterial ECs and gCap ECs within the lungs. **E**, Increase of arterial ECs in distal vascular bed in PH mice. RNASCOPE analysis showing increase of Cxcl12^+^/CD31^+^ ECs in the distal capillary region of CKO mice. n=5 per group. **F** and **G**, A representative UMAP plot showing the lung EC subpopulations change with accumulation of arterial ECs in IPAH patients. **H**, *Egln1*^*Cdh5-CreERT2*^ (iCKO) mice develops spontaneous PH with increase of RVSP and RV hypertrophy. n=3 (WT) or 8 (iCKO). **I**, Cxcl12-DsRed reporter studies showed that Cxcl12 co-localizes with gCap ECs but not aCap ECs in PH mice. More than 3 mice were examined. **J**, Pseudotime trajectory analysis suggested that arterial ECs might be derived from gCap ECs analyzed by Monocle3. **K**, Plvap-DreER^T2^ lineage-tracing study demonstrated that gCap ECs give rise to Cxcl12^+^ arterial ECs in the distal microvascular bed under PH condition. gCap^R^ mice were treated with tamoxifen to label gCap ECs with Tdtomato, followed by SuHx treatment to induce PH development. Cxcl12 was labeled by RNASCOPE. Arrows point to the Cxcl12^+^/Tdtomato^+^ cells, indicative of ArtECs. **L**, Quantification of arterial ECs and gCap derived arterial ECs in PH. n=4 per group. **M**, Transcription factors prediction of lung endothelial subpopulation via SCENIC. SOX17 is one of the top candidates governing arterial EC gene signature. **N**, Pseudotime trajectory showing SOX17 expression across EC subpopulations. Lighter color indicates higher gene expression. **O** and **P**, Sox17 and arterial reprogramming is depended on HIF-2α. A representative UMAP demonstrating HIF-2α deletion (EH2) blocks arterial reprogramming related genes in *CKO* mice by scRNA-seq analysis. **Q**. Western blotting confirmed that Sox17 and Nicd4 expression was downregulated by HIF-2α deletion in CKO mice. **R**, Notch4 neutralized antibodies inhibited RVSP in iCKO mice. n=12 (Anti-IgG) or 13 (Anti-Notch4, 10mg/kg, intraperitoneal injection twice weekly for 2 weeks). **S**, Notch4 neutralized antibodies inhibited RV hypertrophy in iCKO mice. **T**, Notch4 neutralized antibodies reduced arterial gene expression. **U**, A diagram showing the hypothesis of this study. Student t analysis. \*\**p*<0.01. \*\*\**p*<0.001, \*\*\*\**p*<0.0001. Scale bar: 50_µ_m.

Cxcl12 is absent in distal capillary ECs in the lung^4^. Our data demonstrated that Cxcl12 is one of the top arterial EC markers in PH. Thus, we hypothesize that Cxcl12 could be a novel marker for arterial ECs in the microvasculature during PH development. We then bred Cxcl12^DsRed^ reporter mice into *Egln1*^*Cdh5CreERT2*^ mice (iCKO) to generate iCKO;Cxcl12^DsRed/+^ (iCKO^R^) mice. iCKO^R^ mice developed spontaneously PH after tamoxifen treatment (**Fig. H**). There is minimal DsRed^+^ cells in the capillary regions of WT^R^ lungs. DsRed^+^ cells were dramatically increased in the iCKO^R^ lungs and colocalized with gCap ECs marker *Gpihbp1* but not aCap marker *Ednrb* (**Fig. I**). Leveraging the scRNA-seq dataset, our data showed that arterial ECs are likely derived from gCap ECs supported by pseudotime analysis of the lung ECs using Monocle3 (**Fig. J**). The gold-standard strategy to study cell lineage conversion *in vivo* uses genetic lineage tracing. Previous and our scRNA-seq analysis showed that Plvap is a selective marker for gCap ECs^5^. We then generated Plvap-DreER^T2^;CAG-RSR-Tdtomato (gCap^R^) mice and challenged with Sugen5416 (20mg/kg, weekly) and 10% O_2_ hypoxia treatment for 3 weeks (SuHx) to induce PH development. After tamoxifen treatment, Plvap^+^ ECs were successfully labeled with Tdtomato. SuHx challenged gCap^R^ mice showed an increase in arterial ECs (Cxcl12^+^) and gCap-derived arterial ECs (*Cxcl12*^+^/*Tdtomato*^+^) cells in the microvasculature (**Fig. K and L**), suggesting gCap ECs give rise to arterial ECs during PH development.

Multiple molecules such as Sox17 and Notch are involved in activation of arterial specification. SCENIC analysis predicted that Sox17 is one of the top candidates governing arterial EC identity in PH (**Fig. M**). Pseudotime trajectory analysis also show that Sox17 is highly expressed in arterial ECs (**Fig. N**). Through analyzing scRNA-seq data from WT, CKO mice, and *Egln1*^*Tie2Cre*^*/Hif2a*^*Tie2Cre*^ (EH2) double knockout mice, we demonstrated that arterial and capillary EC alterations were normalized (**Fig. O and P**) and Sox17 expression was reduced in EH2 lungs (**Fig. Q**). SOX17 acts as the upstream regulator of Notch signaling, and is required for acquisition and maintenance of arterial identity. Based on our scRNA-seq data, Notch4 activation (Nicd4 expression) is increased in CKO mice (**Fig. Q**). Genetic deletion of HIF-2α normalized Nicd4 expression, suggesting that Notch4 is the HIF-2α/Sox17 downstream target involved in the arterial reprogramming. We then found that neutralized Notch4 signaling inhibited PH development by reducing RVSP and RV hypertrophy, arterial gene expression in iCKO mice compared to IgG control (**Fig. R-T**).

Overall, we demonstrated an increase in arterial ECs in the distal PH lungs. Our results uncovered a novel role of gCap to arterial ECs transition during the pathogenesis of PH development via HIF-2α/Notch4 signaling. Inhibition of Notch4 signaling is a promising avenue to inhibit aberrant accumulation of arterial ECs and prevent pathological gCap-to-arterial reprogramming in PH (**Fig. U**).

## Acknowledgments

All animal protocols were approved by the Institutional Animal Care and Use Committee of University of Arizona. The authors thank the Pulmonary Hypertension Breakthrough Initiative (PHBI) for providing the lung tissues. Funding for the PHBI is provided under an NHLBI R24 grant (R24HL123767). The data, analytical methods, and materials that support the findings of this study will be available to other researchers from the corresponding authors on reasonable request.

