## supplemental material for "General Capillary Endothelial Cells Undergo Reprogramming into Arterial Endothelial Cells in Pulmonary Hypertension through HIF-2α/Notch4 Pathway"

**Methods**

**Human samples**

The use of archived human lung tissues was granted by the UA Institutional Review Board**.** Human IPAH patients and failed donors (control)’ archived lung tissues were obtained from the Pulmonary Hypertension Breakthrough Initiative (PHBI). A table summarizing clinical and demographic characteristics of IPAH patients and control are provided in **Supplemental Table 1**.

**Animals and Experimental Protocol**

All the mice used in this study are C57BL/6J background. *Egln1^Tie2Cre^* (CKO) and *Egln1/Hif2a^Tie2Cre^* (EH2) mice were generated as described previously^1^. *Egln1^f/f^* (WT) mice were bred with *Cdh5CreERT2* mice (obtained from Dr. Ralf Adams, Max Planck Institute for Molecular Biomedicine, Germany) to generate *Egln1^Cdh5CreERT2^* (iCKO) mice. Both male and female WT and iCKO mice at the age of 7 weeks were treated with tamoxifen (20mg/kg) daily for 5 days and rested for ~3 months for PH related phenotypes measurement, tissue collection, and single-cell RNA sequencing analysis.

Plvap-DreER^T2^ mice was obtained from from Shanghai Model Organisms Center, Inc. CAG-rox-Stop-rox(RSR)-Tdtomato mice^2^ were gifted from Dr. Gary Owen at the University of Virginia. Plvap-DreER^T2^ mice were bred with CAG-rox-Stop-rox-Tdtomato mice to Plvap-DreER^T2^;CAG-RSR-Tdtomato (gCap^R^) mice. gCap^R^ mice at the age of 7 weeks were treated with tamoxifen (20mg/kg) daily for 5 days. Three weeks later, these mice were randomly assigned to normoxia or Sugen5416 treatment (20 mg/kg weekly for 3 weeks) and hypoxia (10% O_2_) for 3 weeks.

Cxcl12^DsRed^ reporter mice (JAX: 022458) was bred into the background of iCKO mice to generate iCKO;Cxcl12^DsRed/+^ (iCKO^R^) and WT;Cxcl12^DsRed^ (WT^R^) mice. Both male and female WT^R^ and iCKO^R^ mice at the age of 7 weeks were treated with tamoxifen (20mg/kg) daily for 5 days and rested for ~3 months for PH related phenotypes measurement and tissue collection.

For antibodies treatment, iCKO mice after tamoxifen treatment were treated with 10mg/kg Anti-Notch4 antibodies (InVivoMAb, Cat#BE0129) or anti-IgG (InVivoMAb, Cat# BE0091) intraperitoneally twice weekly for 2 weeks.

**Hemodynamic measurement**

Right ventricular systolic pressure (RVSP) was measured with a 1.4F pressure transducer catheter (Millar Instruments) and recorded with AcqKnowledge software (Biopac Systems Inc.) as described previously. Briefly, the catheter was inserted into the right ventricle in mice under anesthesia (100 mg ketamine/ 5mg xylazine /kg body weight, i.p.)^3^.

**Single-cell RNA sequencing analysis**

We collected whole lung tissues from WT, CKO, and EH2 mice at the age of 3 months for scRNA-seq analysis. Briefly, lung tissues were minced into small pieces and digested with Liberase (Millipore Sigma) for single cell isolation. Equal number of single cells isolated from WT or CKO or EH2 mice was pooled and loaded on 10X Genomics Chromium Single Cell Controller to generate barcoded single cells for construction of single cell cDNA libraries. ScRNA-seq was performed on a Hiseq 4000 with pair-end 150bp (Novogene). The scRNA-seq data were processed using the Cell Ranger pipeline to align reads and generate count matrix. ScRNA-seq data were prefiltered to remove cell with negative hashing tag or doublets and high mitochondrial transcripts (> 10%) employing Seurat v4^4^. Lung cell populations including ECs (*Emcn*, *Pecam1*, *Cdh5* and *Flt1*), SMCs (*Acta2*, *Myh11*, *Cnn1* and *Tagln*), fibroblasts (*Col1a1*, *Col1a2* and *Fn1*), pericytes (*Pdgfrb*, *Notch3 and Cspg4*), alveolar type 1 (*Aqp5*, *Ager* and *Pdpn*) and type 2 (*Sftpb*, *Sftpc* and *Abca3*) epithelial cells, club/ciliated cells (*Scgb1a1*, *Scgb3a2*, *Foxj1* and *Dnali1*), macrophages (*Cd68*, *Cd14*, *Spi1* and *Fcgr2b*), T lymphocytes (*Cd3g*, *Cd3e* and *Cd7*), B lymphocytes (*Cd19* and *Cd22*), lymphatic ECs (*Prox1*, *Flt4* and *Lyve1*) were predicted by cell type-selective markers using UCell package^5^. Lung ECs transcriptomes were extracted for differential gene expression, EC subpopulation clusters, and pseudotime trajectory analysis. For lung EC subpopulations clustering, we defined EC subpopulation according to their markers according previous publication^6,7^. To predict the cell status change, Monocle3 pipeline was performed^8^. Gene regulatory network analysis and transcription factors prediction were performed using the SCENIC tool^9^. For the human scRNA-seq dataset analysis^10^, lung cell populations were annotated and ECs were extracted for subpopulation clusters using Seurat V4 as described above.

**Spatial transcriptomics and analysis**

Mouse lung tissues were perfused with PBS and fixed with 10% formalin via tracheal instillation at a constant pressure (15 cm H_2_O) and embedded in paraffin wax. Lung tissues were sectioned into 5 μm sections. Tissue sections were placed within the fiducial frame or the etched frames of the Capture Area on the 10X Genomics Visium Spatial slides. Slides were then deparaffinated, decrosslinked and stained with H & E staining kit (Millipore Sigma). Images were acquired under Keyence BZ-X800E slide scanner. The mouse whole transcriptome probe panel is added to the deparaffinized, stained, and decrosslinked tissues. After hybridization, single stranded ligation products were released and then captured on the Visium slides. Probes are extended by the addition of UMI, Spatial Barcode and partial Read 1, followed by library preparation and sample indexed. The library was sequenced on a Hiseq 4000 with pair-end 150bp (Novogene). The raw sequencing data was analyzed by CellRanger 7.0 (10X Genomics) and Seurat V4. Visium data was integrated with scRNA-seq data. The cell annotation was transferred from scRNA-seq.

**Immunostaining**

For immunofluorescent staining on these fresh frozen tissue, lung sections (5 μm) were fixed with 4% paraformaldehyde and blocked with 0.1% Triton X-100 and 5% normal goat serum at room temperature for 1 hour. After 3 washes with PBS, the slides were incubated with and anti-SLC6A4 (Thermo Fisher Scientific, Cat# 702076, 1:50) or NOTCH4 (Cell Signaling, Cat#2423, 1:50) at 4°C overnight then incubated with Alexa 594 -conjugated anti-rat or anti-mouse or anti-rabbit IgG (Thermo Fisher Scientific) at room temperature for 1 h. Nuclei were counterstained with DAPI mounting medium (SouthernBiotech, Birmingham, AL, USA).

**RNASCOPE In Situ Hybridization Assay**

A Multiplex Fluorescent V2 RNAscope in situ hybridization assay (Advanced Cell Diagnostics, Newark, CA) were performed on lung cryosections from IPAH patients and failed donor subjects, and mice lung samples. Briefly, the tissue sections were fixed in chilled 10% neutral buffered formalin for 1h at 4°C, followed by dehydrated with ethanol. Then the sections were incubated with hydrogen peroxide at RT for 10 mins, followed by Protease IV incubation for 30 mins at RT. The sections were hybridized with different probes [Advanced Cell Diagnostics , human probes: NOTCH4 (409631-C2), EDNRB (528301-C2); mouse probes, Cxcl12 (422711-C1), Ednrb(473801-C2), Gpihbp1 (540631-C3); and Tdtomoto (317041-C1)] for 2 h at 40°C followed by signal amplification for 30 minutes using RNAscope® Multiplex Fluorescent v2 Assay (Advanced Cell Diagnostics) as per manufacturer’s instructions. The signal was developed by incubating the slides with HRP-C1 for 15 minutes, and Opal Dye 620 for 30 mins, followed by incubation with HRP-C2 or C3 for 15 minutes and Opal Dye 690 for 30 mins. Then the slides were washed with PBS, counterstained with DAPI, mounted in DAPI mounting medium (SouthernBiotech, Birmingham, AL, USA) and images were captured.

**Western blot**

Protein lysates were prepared by mincing lung tissues from mice in cold NP-40 lysis buffer supplemented with protease inhibitor cocktails (MilliporeSigma). Equal amount of protein was loaded for SDS-PAGE and Western Blotting. The PVDF membranes were blotted with anti-NOTCH4 (Cell Signaling Technology, Cat#2423, 1:1000), anti-SOX17 (Abcam, #ab224637, 1:10,000), anti-CXCL12 (Santa Cruz Biotechnology, Cat# sc-74271, 1:500) or anti-β-actin (Sigma-Aldrich, #A2228) antibodies.

**Data availability**

Scripts used for scRNA sequencing and Visium analysis in R objects are available in GitHub (https://github.com/DaiZYlab/gCap-to-artEC). Other data that support the findings of this study are available from the corresponding author upon reasonable request.

**Statistical Analysis**

Statistical determination was performed on Prism 9 (Graphpad Software Inc.). Two-group comparisons were compared by the unpaired 2-tailed Student t test for equal variance or the Welch t test for unequal variance. Multiple comparisons were performed by one-way ANOVA with a Tukey post hoc analysis that calculates corrected P values. P less than 0.05 indicated a statistically significant difference. All bars in dot plot figures represent mean. All bar graphs represent mean±SD.

**Acknowledgements**

The authors thank the Pulmonary Hypertension Breakthrough Initiative (PHBI) for providing the Data/tissue samples. Funding for the PHBI is provided under an NHLBI R24 grant (R24HL123767) and by the Cardiovascular Medical Research and Education Fund. This work was supported in part by NIH grant R00HL138278, R01HL158596, R01HL62794, R01HL169509, R01HL170096, AHA Career Development Award 20CDA35310084, The Cardiovascular Research and Education Foundation, Arizona Biomedical Research Centre funding (RFGA2022-01-06), and University of Arizona startup funding to Z.D.

**Author contributions**

Z.D. conceived the experiments and interpreted the data. B.L., D.Y., X.X., K.R., H.Z., Y. C., R.D., A.G., H.D., and Z.D. designed, performed experiments, and analyzed the data. Z.D. wrote the manuscript. S.Q., V.K., M.F., revised the manuscript.

**Supplemental Table**

**Supplemental Figure**

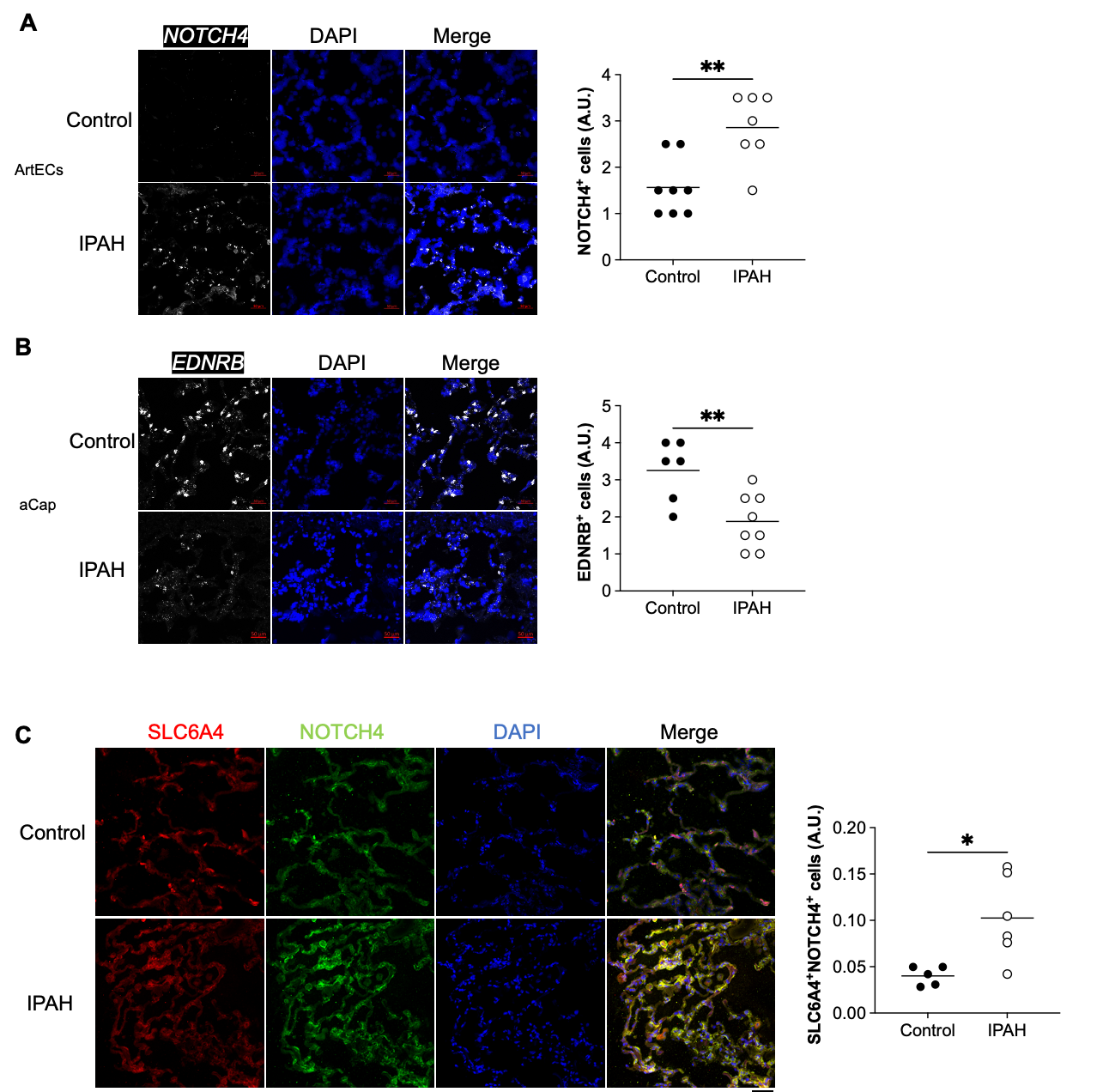
 **Supplemental Figure 1, Arterial ECs were increased and aCap ECs were reduced in the lung of IPAH patients.**

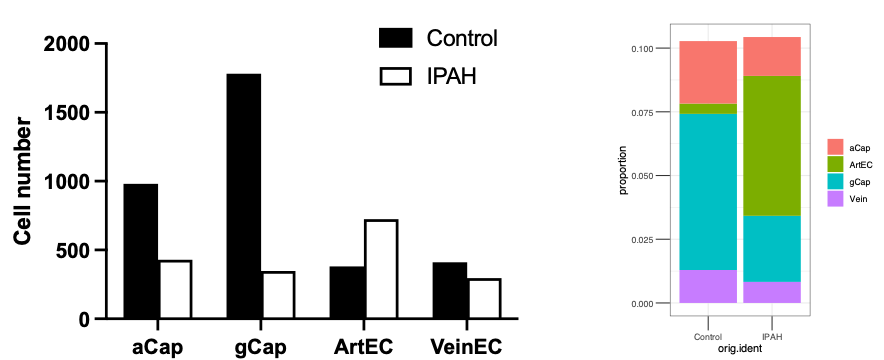

**Supplemental figure 2, Cell number and proportional analysis of EC subpopulations in human IPAH lungs.**

| **PAH or Failed Donor** | **Gender** | **Race** | **Age** |
| --- | --- | --- | --- |
| IPAH1 | Female | White | 14 |
| IPAH2 | Female | White | 11 |
| IPAH3 | Female | White | 39 |
| IPAH4 | Female | White | 16 |
| IPAH5 | Male | White | 53 |
| IPAH6 | Male | White | 13 |
| IPAH7 | Male | Asian | 29 |
| IPAH8 | Male | Asian | 18 |
| IPAH9 | Female | White | 55 |
| IPAH10 | Female | White | 57 |
| Failed donor 1 | Female | White | 49 |
| Failed donor 2 | Female | White | 57 |
| Failed donor 3 | Male | White | 49 |
| Failed donor 4 | Male | White | 49 |
| Failed donor 5 | Female | White | 43 |
| Failed donor 6 | Male | White | 30 |
| Failed donor 7 | Female | Hispanic | 55 |
| Failed donor 8 | Male | Unknown | 13 |
| Failed donor 9 | Female | Asian | 34 |
| Failed donor 10 | Male | White | 21 |

**Extended Data Table 3, Clinical and demographic characteristics of PAH patients and failed donors.**

**Extended Data Fig. 1, Increased arterial ECs and decreased aCap ECs in PH mice.** a, Immunostaining and RNASCOPE analysis showing CKO mice exhibited upregulation of arterial EC marker Sox17 and downregulation of aCap EC marker *Ednrb* compared with WT mice. Student t analysis. **p*<0.05, ****p*<0.001.

**Extended Data Fig. 2, *Egln1^Cdh5CreERT2^* (iCKO) mice developed spontaneously PH and arterial programming. a,** Hemodynamic measurement showing that iCKO mice had increased right ventricular systolic pressure (RVSP) compared with *Egln1^f/f^* (WT) mice. **b**, Cardiac dissection showed the upregulation of right heart and left heart hypertrophy in iCKO mice compared with WT mice. **c,** A representative UMAP plot showing the abnormal EC subpopulations change with accumulation of arterial ECs (green) in iCKO mice at the age of 5 months. **d**, Cell proportion change of EC subpopulations in **c**. **e**, A representative heatmap showing the top markers selectively enriched for each EC cluster, the data was generated by integrated data from WT and iCKO. Student t analysis. *****p*<0.0001.

**Extended Data Fig. 3, Spatial transcriptomics analysis showing the cellular change in CKO mice. Pan** EC markers (Pecam1 and Cdh5) and EC subpopulations including arterial ECs (Cxcl12 and Sparcl1, a), aCap ECs (Car4 and Ednrb, b), and gCap ECs (Gpihbp1 and Plvap, c) were identified within tissue sections from WT mice and CKO mice.

**Extended Data Fig. 4. scRNA-seq analysis on lung tissues from IPAH patients. a**, A representative heatmap showing the top markers selectively enriched for each EC cluster, the data was generated by integrated data from human IPAH patients and failed donors. **b**, the overlapped genes between human pulmonary arterial ECs and mouse pulmonary arterial ECs.

**Extended Data Fig. 5. Increase of arterial endothelial cells and loss of aCap endothelial cells in IPAH lungs**. RNASCOPE analysis showing upregulation of arterial EC marker NOTCH4 (a) and downregulation of aCap EC marker *EDNRB* (b) in IPAH patients compared with control donors.

**Extended Data Fig. 6, Cxcl12 is an arterial EC marker. a,** a diagram showing the strategy of generating Cxcl12 reporter in iCKO mice background**.** iCKO;Cxcl12^DsRed/+^ (iCKO^R^) mice developed spontaneously PH evident by increased RVSP (**b**) and RV hypertrophy (**c**). **d**, Increase of arterial ECs and loss of aCap ECs in iCKO^R^ mice. RNASCOPE assay was performed to label gCap and aCap ECs. Student t analysis. ***p*<0.01, ****p*<0.001.

**Extended Data Fig. 7, Plvap-DreER^T2^ labeled general capillary ECs in mice.** RNASCOPE assay demonstrated that Plvap-DreER^T2^ labeled general capillary (gCap) ECs but not aerocyte (aCap) ECs.

**Extended Data Fig. 8, aCap ECs were reduced in Sugen5416/hypoxia challenged mice.** RVSP and RV hypertrophy were increased in SuHx challenged Plvap-DreER^T2^ mice compared to basal mice. Student t analysis. **p*<0.05, ***p*<0.01, ****p*<0.001.

**Extended Data Fig. 9, Molecular alteration of arterial ECs in PH mice.** **a**, A heatmap showing many genes related to arterial EC markers, angiocrine factors, and glycolysis were upregulated in the arterial ECs of CKO mice. **b**, A Dotplot showing the increased ligand-receptor interaction between arterial ECs and other perivascular cell types including fibroblasts, smooth muscle cells and pericytes in CKO mice compared with WT mice.

**Extended Data Fig. 10, HIF-2/Notch4 signaling controlled arterial programming in PH mice.** **a**, Bulk RNA-seq analysis showed that HIF-2α deletion inhibited arterial gene expression in CKO mice. **b**, Western blot analysis demonstrated that Notch4 but Notch1 is activated in the lungs of CKO mice. c, Notch4 activation is depended on HIF-2α. HIF-2α deletion in CKO mice inhibited Notch4 activation assessed by Nicd4 expression. Student t analysis (**b**). ANOVA with Tukey post hoc analysis (**c**). Student t analysis, **p*<0.05.
